# Mediobasal hypothalamic FKBP51 acts as a molecular switch linking autophagy to whole-body metabolism

**DOI:** 10.1101/2021.05.31.445775

**Authors:** Alexander S. Häusl, Lea M. Brix, Thomas Bajaj, Max L. Pöhlmann, Kathrin Hafner, Meri De Angelis, Joachim Nagler, Georgia Balsevich, Karl-Werner Schramm, Patrick Giavalisco, Alon Chen, Mathias V. Schmidt, Nils C. Gassen

**Author notes:** Correspondence: Mathias V. Schmidt or Nils C. Gassen. shared senior authorship (Mathias V. Schmidt, Nils C. Gassen).

## Abstract

The mediobasal hypothalamus (MBH) is the central region in the physiological response to metabolic stress. The FK506-binding protein 51 (FKBP51) is a major modulator of the stress response and has recently emerged as a scaffolder regulating metabolic and autophagy pathways. However, the detailed protein-protein interactions linking FKBP51 to autophagy upon metabolic challenges remain elusive. We performed mass spectrometry-based metabolomics of FKBP51 knockout (KO) cells revealing an increased amino acid and polyamine metabolism. We identified FKBP51 as a central nexus for the recruitment of the LKB1/AMPK complex to WIPI4 and TSC2 to WIPI3, thereby regulating the balance between autophagy and mTOR signaling in response to metabolic challenges. Furthermore, we demonstrated that MBH FKBP51 deletion strongly induces obesity, while its overexpression protects against high-fat diet (HFD) induced obesity. Our study provides an important novel regulatory function of MBH FKBP51 within the stress-adapted autophagy response to metabolic challenges.

## Introduction

An adequate response to nutritional changes requires a well-coordinated interplay between the central nervous system and multiple peripheral organs and tissues to maintain energy homeostasis. High-caloric food intake and chronic overnutrition significantly challenge this system on a cellular and organismic level and are main drivers in the development of obesity, a hallmark of the metabolic syndrome ^1^.

Autophagy is an evolutionarily conserved process that efficiently degrades cellular components, like unfolded proteins or organelles, to provide internal nutrients and building blocks for cellular fitness ^2^. Dysfunctional autophagy is associated with many diseases, such as neurodegeneration, liver disease, cancer, and metabolic syndrome ^3–5^. In obesity, however, the alterations of autophagy are not fully explored yet. Autophagic signaling in obese subjects is suppressed in pancreatic B-cells, liver, and muscle whereas other studies demonstrated enhanced autophagy signaling in adipose tissue ^3,6,7^. Autophagy initiation is tightly controlled by a series of proteins encoded by autophagy-related genes (ATG) and regulatory hetero-protein complexes, including the systemic energy sensor 5’-AMP-activated protein kinase (AMPK) that is in balance with the mechanistic target of rapamycin (mTOR) ^8^. Amino acid surplus activates mTOR signaling, which further suppresses autophagy by inhibiting the UNC51-like kinase 1 (ULK1) complex (comprised of ULK1, ATG13, FIP200, and ATG17). In the course of nutrient deprivation, elevated AMP actives AMPK to initiate autophagy via the ULK1 complex, and in turn diminishes mTOR signaling ^9,10^. It has recently been shown that the WD-repeat proteins that interact with the phosphoinositides protein family (WIPI proteins) act as subordinate scaffolders of the LKB1/AMPK/TSC2/FIP200 network linking AMPK and mTOR signaling to the control of autophagy upon metabolic stress ^11^.

In the mediobasal hypothalamus (MBH), the brain’s central region for metabolic control, the deletion of ATG7 (a ubiquitin E1-like ligase, downstream of the ULK1 complex) in proopiomelanocortin (POMC)-expressing neurons resulted in obesity and dampened sympathetic outflow to white adipose tissue (WAT) ^12^, while ATG7 deficiency exclusively in agouti-related protein (AgRP)-expressing neurons resulted in decreased body weight ^13^. Together, this data indicates a role of MBH autophagy in the development of obesity, but key regulatory proteins remain largely elusive.

The FK506-binding protein 51 (FKBP51, encoded by *Fkbp5*) is the main modulator of the stress response and is best characterized as a co-chaperone to HSP90, thereby orchestrating diverse pathways important to maintain homeostatic control ^14–17^. We and others have provided evidence that FKBP51 is associated with type 2 diabetes and have shed light on its role as a fundamental regulator of obesity and glucose metabolism ^18–22^. Following the identification of FKBP51 as a negative regulator of the serine/threonine kinase AKT in cancer cells ^23^, we showed that FKBP51 acts as a modulator of glucose uptake by mediating the AKT2/PHLPP/AS160 complex specifically in muscle ^18^. Furthermore, we demonstrated that FKBP51 induces autophagy through autophagy-promoting BECN1 in two ways: (1) FKBP51 limits AKT-directed inhibitory phosphorylation of BECN1 at S234 and S295 ^24,25^, and (2) it reduces AKT-mediated phosphorylation of SKP2 at S72 and thereby lowering its E3-ligase activity preventing BECN1 from proteasomal degradation ^26^. Autophagy and FKBP51 are involved in the regulatory role of adipocyte differentiation and mass development ^19,27–29^ and are upregulated in the MBH following starvation ^13,30,31^. The data indicate converging mechanisms of FKBP51-directed protein scaffolding and autophagy and therefore we hypothesized a subordinate role of FKBP51 in concert with members of the autophagy signaling network in shaping central and peripheral autophagy signaling. In the current study, we set out to identify the molecular interplay of FKBP51 with cellular autophagic signaling and tested whether FKBP51 shapes the *in vivo* whole-body response to an obesogenic challenge. Our study unravels the tissue specificity of autophagy signaling in response to obesity and uncovers FKBP51 as a novel regulatory link between the stress-induced LKB1/AMPK-mediated autophagy induction and WIPI protein scaffolds.

## Results

### FKBP51 deletion increases AMPK and mTOR-associated amino acid and polyamine biosynthesis

As a first approach to broaden our insights into the contribution of FKBP51 under basal (1x glucose) and metabolically challenging (2x glucose) conditions, we performed a multilevel mass spectrometry-based metabolomics profiling analysis of human neuroblastoma SH-SY5Y cells lacking FKBP51 (FKBP51 KO) and corresponding wildtype controls (WT). FKBP51 KO cells showed a significant increase in multiple metabolites compared to WT cells to different extents under basal and high glucose conditions (Fig. S1A – S1D). Most frequent and pronounced alterations could be attributed to biosynthetic and metabolic pathways of various amino acids, including but not limited to arginine, valine, leucine, and isoleucine biosynthesis and histidine, cysteine, and methionine metabolism (Fig. 1A, Fig. S1E).

**Fig. 1:**
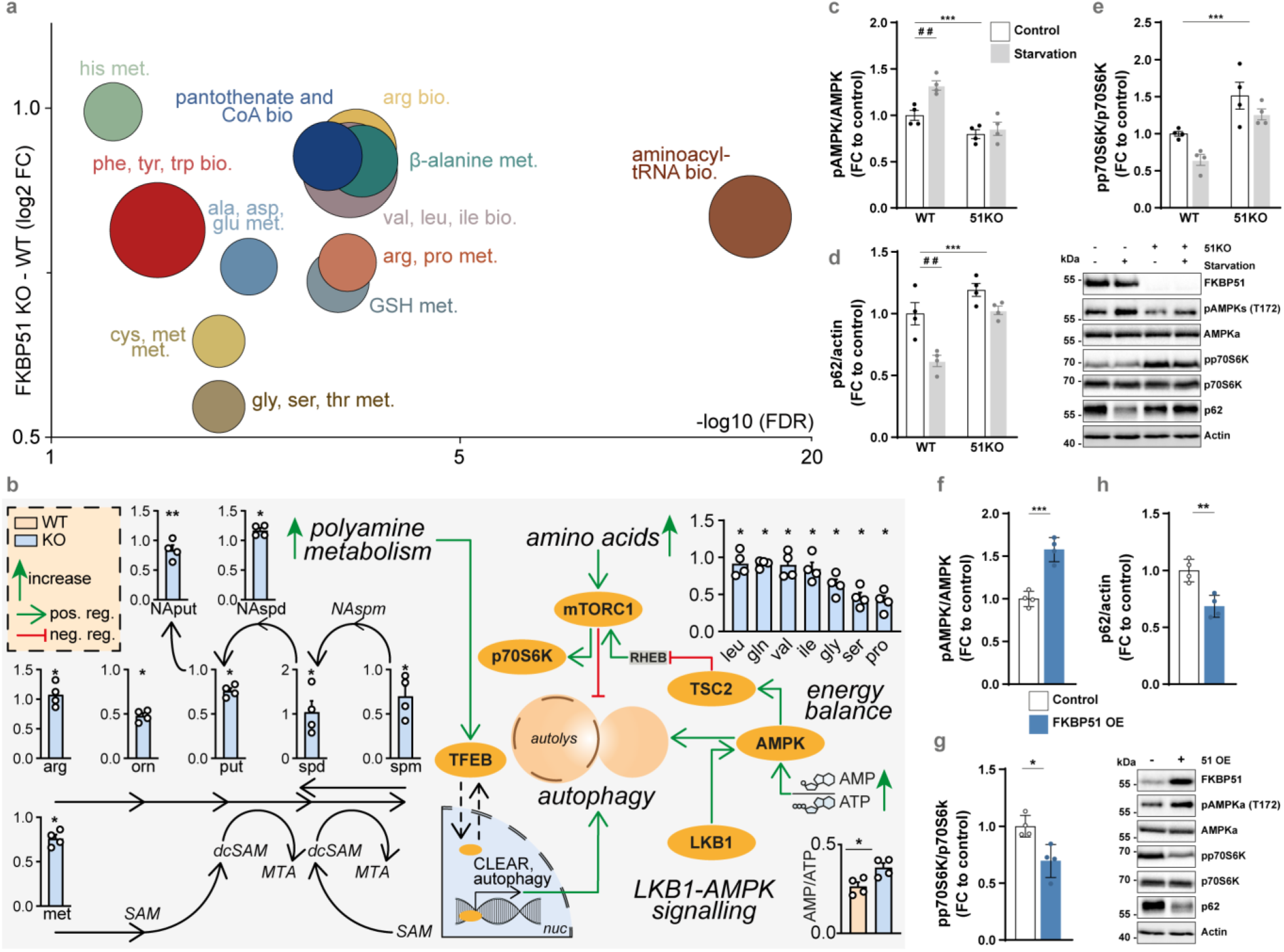
FKBP51 regulates AMPK and mTOR activity as well as associated amino- and polyamine biosynthesis pathways. **(A)** Analysis and regulation of significantly altered pathways of FKBP51 knockout (KO) and wildtype (WT) cells. The f(x)-axis shows the (median) log2 fold change (FC) of all significantly altered metabolites of the indicated pathway and the false discovery rate (FDR, equals the –log10 adjusted p-value) is shown on the x-axis. The size of the circles represents the amount of significantly changed metabolites in comparison to all metabolites of a particular pathway. **(B)** FKBP51 deletion increases metabolites of the polyamine pathway, the AMP/ATP ratio and enhances levels of amino acids associated with mTOR signaling. **(C)** WT or FKBP51 KO cells were starved in HBSS medium for 4 h to induce autophagy and the quantification of pAMPK (T172), **(D)** p62, and **(E)** pp70S6K (T389). FKBP51 overexpression (FKBP51 OE) in Neuro2a cells (see Suppl. Fig. 2D for validation) enhanced autophagy signaling. Quantification of **(F)** pAMPK (T172), **(G)** pp70S6K (T389), and **(H)** p62. All data for **(C-H)** are shown as relative fold change compared to control condition; ± s.e.m.; * p < 0.05, **p < 0.01, ***p < 0.001; # p < 0.05, ## p < 0.01. Two-way ANOVA was performed in **(C-E)** and followed by a Tukey’s multiple comparisons test. The unpaired student’s t-test was performed for **(F-H)**. * = significant treatment effect; # = significant genotype effect.

It is well known that amino acids signal to mTOR and that mTOR itself actively participates in the sensing of amino acids in the lysosomal lumen. Particularly, the branched-chain amino acids (BCAAs) leucine, valine, and isoleucine, all elevated in FKBP51 KO cells (Fig. 1B, upper right part), can activate mTOR and thereby block the autophagy pathway ^32–34^. On the other hand, the pathway analysis showed increased AMP/ATP ratio and strongly increased levels of polyamines and their metabolites in FKBP51 KO cells (Fig. 1B, left part and lower right part). This indicates increased AMPK activation ^35^ and enhanced hypusination of the translation factor elF5A and thereby positively regulates autophagy via expression of TFEB ^36^. The overall increase in amino acids most likely results from ubiquitous protein degradation, in line with enhanced cellular catabolic processes. We performed isotope tracing with ^13^C-labeled glucose to determine the intracellular flux in FKBP51 KO cells. Here, we observed no substantial differences between both cell types (Fig. S1F) corroborating the fact that cellular catabolic processes rather dominate over anabolic processes. Given the observed effects of FKBP51 deletion on polyamine and amino acid biosynthesis, we hypothesized that FKBP51 affects autophagy and mTOR signaling and were further encouraged to disentangle the underlining metabolic pathways.

### FKBP51 is essential for homeostatic autophagy following nutrient deprivation

Since autophagy and FKBP51 expression levels are highly induced after starvation ^4,31^, we tested whether deletion of FKBP51, in turn, affects the induction of autophagy after nutrient deprivation. To do so, we exposed FKBP51 KO or WT cells to HBSS (Hank’s balanced salt solution) for 4 h to induce cellular starvation.

FKBP51 KO cells showed less phosphorylated AMPK at T172 and lower levels of LKB1 compared to WT cells already under nutrient-rich conditions (Fig. 1C, Fig. S2A), while there was a significant increase in activating phosphorylation of SKP2 at S72 and AKT at S473 (Fig. S2B, S2C). These changes in upstream signaling resulted in the slight but not significant accumulation of the autophagy receptor and substrate p62 (Fig. 1D), which is an important measure to determine autophagic activity ^37^. On the other hand, we could observe increased levels of pp70S6K at T389 in FKBP51 KO cells (Fig. 1E). These data imply a reduction of autophagy signaling and increased mTOR signaling after FKBP51 deletion under basal conditions ^38^. Importantly, the deletion of FKBP51 blocked the starvation-induced increase in LKB1 protein as well as the activation of AMPK at T172 (Fig. 1C, Fig. S2A) and thereby significantly reduced the level of autophagy signaling (Fig. 1D). These data underline the importance of FKBP51 in the autophagic stress response after starvation and suggest a tight regulation of FKBP51 on AMPK and LKB1.

Next, we investigated whether FKBP51 upregulation can enhance autophagy by moderately overexpressing (OE) Flag-tagged FKBP51 in Neuro2a cells (Fig. S2D). In line with our hypothesis, FKBP51 OE resulted in highly increased phosphorylation of AMPK at T172 and enhanced levels of LKB1 (Fig. 1F, Fig. S2E) which indicates an increase of upstream autophagy initiation, further evidenced by increased phosphorylation of ULK1 at S555 (Fig. S2E). Consequently, there was an increase in pro-autophagic phosphorylation of BECN1 at S14 and S91/S94 (in humans at S93/S96) through kinases ULK1 and AMPK, respectively (Fig. S2E) ^39^. Previous literature has shown that increased LKB1/AMPK signaling also activates the TSC1/TSC2 complex, which in turn inhibits mTOR activity ^40^. In line with that, phosphorylation of TSC2 at S1387 was significantly enhanced (Fig. S2E) and the phosphorylation of AKT at S473, SKP2 at S72 and the mTOR substrate pp70S6K at T389 were significantly decreased (Fig. 1G, Fig. S2F). Finally, we validated autophagy signaling by the increase in phosphorylation of ATG16L at S278, a novel marker for autophagy induction ^41^, as well as the decrease in the autophagy substrate p62 (Fig. 1H, Fig. S2G).

Together, these findings demonstrate that FKBP51 regulates autophagy induction, especially after a metabolic challenge. However, it is still unclear whether FKBP51 shapes the autophagic response via direct protein-protein interactions to AMPK and LKB1. Given the fact that FKBP51 was previously shown to interact with several signaling molecules within the autophagy pathway (Fig. 2A), we were encouraged to unravel the underlying molecular mechanism and identify potential novel interactions of FKBP51 with members of the autophagy signaling network.

**Figure 2:**
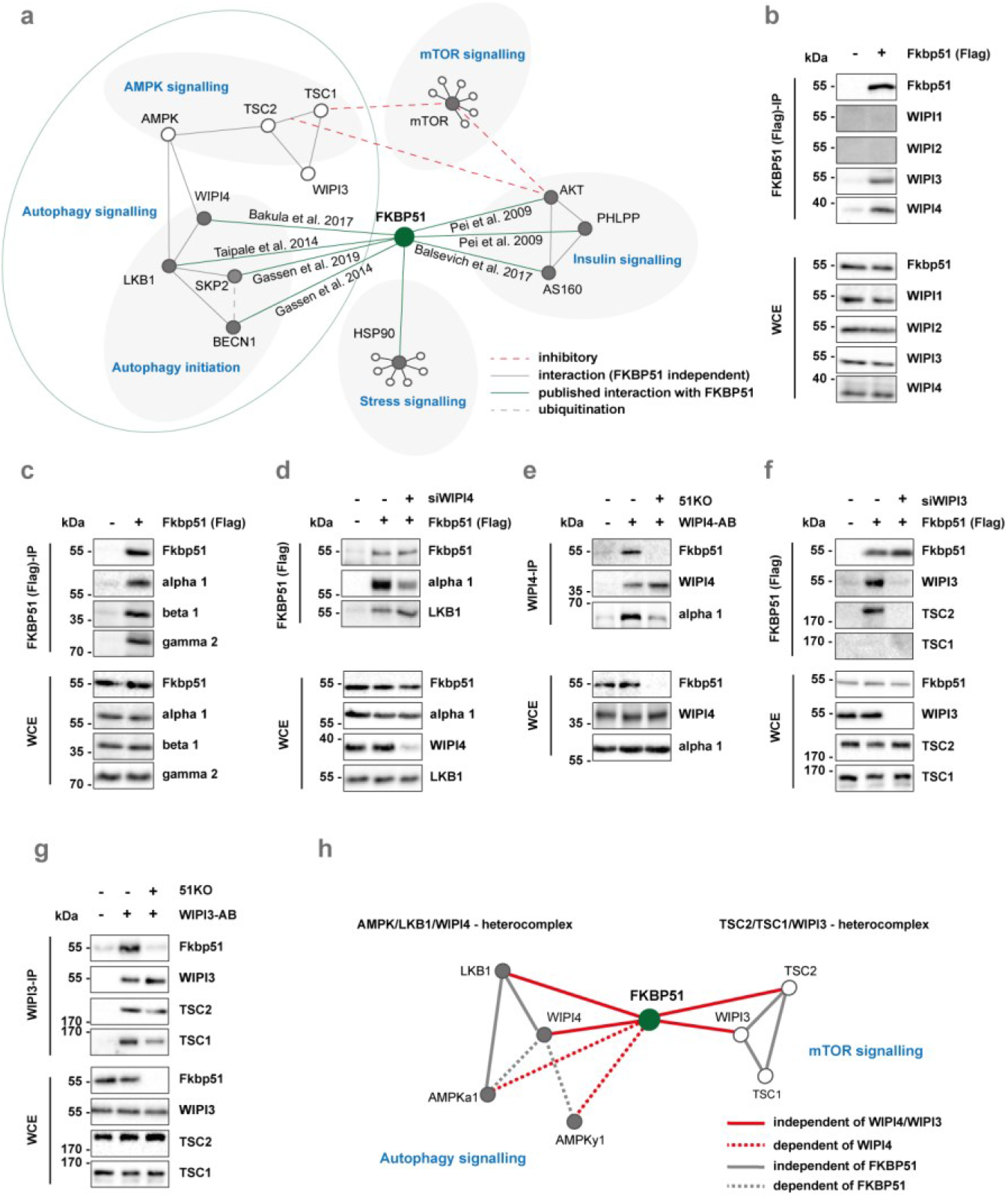
FKBP51 associates with AMPK, TSC2, and WIPI3 & 4 to regulate autophagy and mTOR signaling. **(A)** Published protein-protein interactions of FKBP51. **(B)** FKBP51 associates with WIPI3 and WIPI4, but not with WIPI1 and WIPI2. **(C)** Interaction of FKBP51 with AMPK subunits. **(D)** Interaction of FKBP51 with LKB1 and AMPK in WIPI4 KO cells. **(E)** Interaction of WIPI4 with AMPK in FKBP51 lacking cells. **(F)** FKBP51 interacts with TSC2 in dependency of WIPI3. **(G)** WIPI3 interacts with TSC2 and TSC1 in presence or absence of FKBP51. **(H)** Novel associations of FKBP51 in the regulation of autophagy and mTOR signaling and the proposed model of interaction. FKBP51 recruits LKB1 to the AMPK-WIPI4 complex and thereby facilitates AMPK activation. Furthermore, FKBP51 scaffolds TSC2-WIPI3 binding to alter mTOR signaling.

### FKBP51 associates with LKB1/AMPK/WIPI4, and TSC2/WIPI3 hetero-protein complexes to regulate autophagy and mTOR signaling

A recent study by Bakula and colleagues investigated the role of the four WIPI proteins on autophagy ^11^ and the WIPI protein interactome revealed an association of WIPI4 with FKBP51, AMPKα1, and AMPKγ2 ^11^. WIPI proteins are essential scaffolding proteins that function as central molecular hubs in order to link key regulatory elements of autophagy with proteins that are sensitive to main metabolic cascades such as amino acid and glucose metabolism ^42^. Based on our FKBP51 starvation and OE experiments, we hypothesized that FKBP51 might scaffold WIPI4 and AMPK to induce autophagy initiation and therefore performed co-immunoprecipitation (co-IP) studies using Neuro2a cells, with FKBP51-Flag OE. These experiments confirmed that FKBP51 associates with WIPI4, but not with WIPI1 and WIPI2 (Fig. 2B) as suggested by Bakula and colleagues ^11^. Intriguingly, our experiments revealed a novel association of FKBP51 with WIPI3 (Fig. 2B). Next, we assessed the association of FKBP51 with various isoforms of AMPK and validated the expected interactions of FKBP51 with AMPKα1, AMPKγ2, and further revealed a novel interaction with AMPKβ1 (Fig. 2C).

To investigate the functional relevance of WIPI4 in the association of FKBP51 with AMPK, we generated WIPI4 knockdown (KD) Neuro2a cells using siRNA co-transfected with FKBP51-Flag (Fig. S2H). We observed that FKBP51 binding to AMPK was reduced in cells lacking WIPI4 (Fig. 2D, Figure S2I). Our precipitation studies further demonstrated that FKBP51 interacts with LKB1, which confirmed the finding of a previous interactomics-based screening experiment by Taipale and colleagues ^16^, and further revealed that this interaction is independent of WIPI4 (Fig. 2D, Fig. S2J). Furthermore, we could demonstrate that WIPI4 deletion blocked the phosphorylation effect of FKBP51 OE on pAMPK at T172 (Fig. S2K), suggesting that FKBP51 binds to AMPK in dependency of WIPI4. In addition, studies in FKBP51 KO cells revealed that the WIPI4 interaction to AMPK depends on the presence of FKBP51 (Fig. 2E), indicating a subordinate role of FKBP51 as a molecular bridge that links the autophagy-relevant WIPI-network to AMPK signaling.

Finally, we assessed the functional relevance of WIPI3-FKBP51 binding. Performing co-IP studies in Neuro2a cells in the presence or absence of WIPI3 (Fig. S2L), we observed that FKBP51 interacts with TSC2 but not TSC1 (Fig. 2F), which are upstream master-regulators of mTOR ^43^. Further, we confirmed the association of FKBP51 with TSC2, which depends on the presence of WIPI3 (Fig. 2F), whereas the interaction of WIPI3 to TSC2 and TSC1 is independent of FKBP51 (Fig. 2G).

Collectively, we demonstrate that FKBP51 regulates autophagy in concert with the WIPI protein family, specifically WIPI3 and WIPI4 via interactions with the mTOR and AMPK cascades (Fig. 2H). The combined interpretation of our molecular and metabolomic analyses led us to hypothesize that FKBP51 regulates autophagy and consequently whole-body metabolism *in vivo*, especially after a metabolic challenge.

### MBH FKBP51 is a regulator of body weight and food intake

We and others have previously shown that FKBP51 acts in a tissue-specific manner in soleus muscle (SM), epididymal WAT (eWAT), and the hypothalamus to regulate metabolism ^18,19,22,30^. However, the interconnected response of autophagy and FKBP51 to a metabolic stressor, such as HFD, remains elusive and encouraged us to investigate the possible relationship between FKBP51 expression and autophagic flux (the protein turnover through catabolic autophagy).

Our analysis revealed that C57/Bl6 mice showed significantly increased FKBP51 protein levels (Figure 3A) and diminished accumulation of the autophagy substrate p62 in the MBH upon 10 weeks of HFD (58% kcal from fat) (Figure 3B). HFD *per se* did not affect the lipidation of the autophagosome-spiking protein light-chain 3 (LC3B-I) to LC3B-II (Figure 3C), which binds the autophagosome and is a reliable marker to analyze autophagic flux ^37^. However, levels of LC3B-II were similarly increased in both conditions after treatment with chloroquine (50 mg/kg), an inhibitor of lysosomal acidification and autophagosome-lysosome fusion that in turn blocks degradation of autophagosome cargo ^37^. In line with the reduced levels of p62, these results indicate an active central autophagic flux after HFD.

**Fig. 3:**
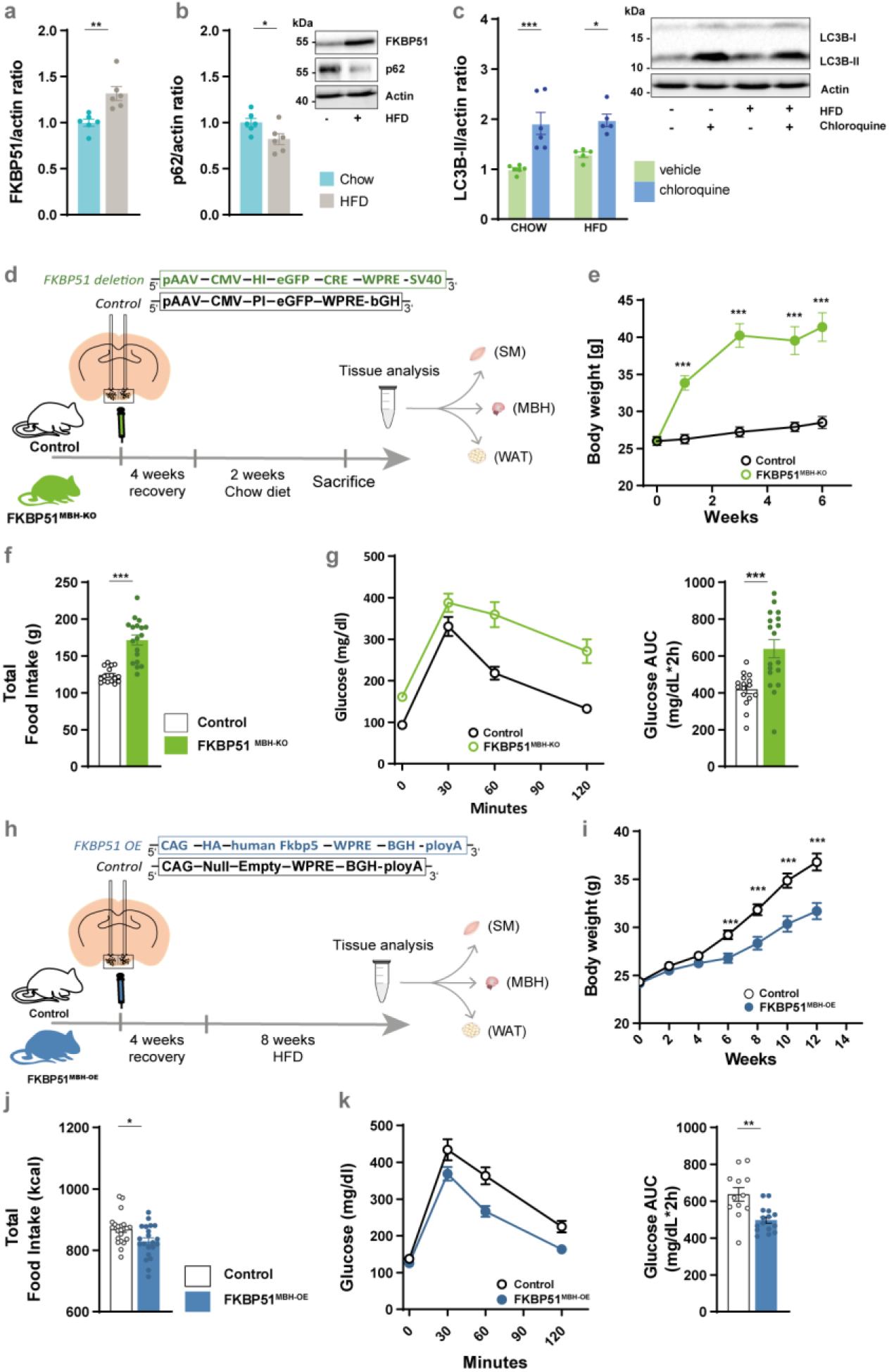
MBH FKBP51 regulates body weight gain, food intake and glucose metabolism. **(A)** 10 weeks of HFD increased hypothalamic FKBP51 in the mediobasal hypothalamus (MBH) (n (chow) = 6 vs. n (HFD) = 6). **(B)** Effects of HFD on the accumulation of p62. **(C)** Treatment with chloroquine (50mg/kg) increased LC3B-II level under chow and HFD conditions. **(D)** FKBP51^lox/lox^ animals were injected with 200nl of *Cre*-expressing virus and fed a chow diet for 6 weeks. **(E)** FKBP51^MBH-KO^ showed significant body weight increase after virus injection on a regular chow diet. **(F)** FKBP51^MBH-KO^ animals showed increased food intake and **(G)** enhanced glucose intolerance. **(H)** For FKBP51 overexpression, animals were injected with an AAV virus into the MBH. **(I)** FKBP51^MBH-OE^ animals showed reduced body weight gain on a HFD diet compared to their control animals **(J)** FKBP51^MBH OE^ animals showed reduced food intake. **(K)** FKBP51^MBH-OE^ animals showed improve glucose tolerance under HFD conditions. For **(A, B, F, G, J, and K)** an unpaired student’s t-test was performed. For **(C)** a two-way ANOVA was performed followed by a Tukey’s multiple comparisons test. For **(E, I)** a repeated measurements ANOVA was performed. ± SEM; * p < 0.05, **p < 0.01, ***p < 0.001.

In peripheral eWAT and SM, however, chloroquine treatment did increase LC3B-II levels only in chow-fed animals (Figure S3A and S3B), implying reduced or even blocked autophagy signaling under HFD conditions. This hypothesis is supported by an increased accumulation of p62 in these peripheral tissues with a significant increase in SM (Figure S3C). Interestingly, FKBP51 protein levels were unaffected in eWAT and slightly but not significantly decreased in SM (Figure S3D). These data suggested a possible interconnected role of FKBP51 and autophagy flux, particularly in the MBH. We, therefore, decided to further test the effects of FKBP51 on autophagy signaling and its significance on whole-body metabolism *in vivo*, by manipulating FKBP51 in the MBH.

First, we injected a *Cre*-expressing virus into the MBH of FKBP51^lox/lox^ animals (Figure 3D) to evaluate the effects of central FKBP51 deletion. Intriguingly, FKBP51^MBH-KO^ animals showed a massive increase in body weight six weeks after surgery, despite their regular chow diet (Figure 3E). The bodyweight increase was accompanied by increased food intake and decreased glucose tolerance (Figure 3F; Figure 3G). These findings were surprising, considering the lean phenotype of full-body FKBP51-deficient mice after prolonged exposure to a HFD ^18,19^ and further highlight the tissue specificity of FKBP51.

Next, we injected an AAV-mediated FKBP51 OE virus into the MBH of C57/Bl6 mice (FKBP51^MBH-OE^, Figure 3H). FKBP51^MBH-OE^ animals showed no substantial differences in body weight gain within the first 4 weeks after surgery. Therefore, we challenged FKBP51^MBH-OE^ animals with a metabolic stressor by feeding them a HFD for eight weeks. Interestingly, animals overexpressing FKBP51 displayed significantly reduced body weight gain compared to the control group (Figure 3I), paralleled by a reduction in food intake (Figure 3J). In a second cohort of FKBP51^MBH-OE^ animals (with an identical body weight phenotype (Figure S3E)), we investigated whether glucose metabolism was altered and observed improved glucose tolerance and insulin sensitivity compared to the control group under HFD, but not under normal chow diet (Figure 3K, Figure S3F - S3H). Together, these experiments reveal an essential role of MBH FKBP51 in central coping mechanisms with an obesogenic stressor and position MBH FKBP51 as a key regulator of whole-body metabolism.

### MBH FKBP51 fine-tunes autophagy signaling in an inversed u-shaped manner

Given the opposing phenotypes of FKBP51^MBH-KO^ and FKBP51^MBH-OE^ animals and the novel regulatory function of FKBP51 in the metabolic control of autophagy, we were interested in the underlying regulation of autophagy signaling. In our KO experiment, viral injection resulted in a high deletion rate within the MBH of FKBP51^MBH-KO^ animal (Figure 4A, 4B). According to our hypothesis, we observed a reduced binding of LKB1 and AMPK to WIPI4 (Figure 4C, see Figure S4A for quantification), which was accompanied by a reduction in the phosphorylation of AMPK at T172, causing less active AMPK (Figure 4D). Downstream of AMPK we monitored diminished phosphorylation of ULK1 at S555, BECN1 at S91/S94 and TSC2 at S1387 (Figure S4B – S4D). Parallel to the effects of AMPK downstream proteins, we detected reduced levels of TSC2 binding to WIPI3 (Figure 4E, see Figure S4E for quantification). Furthermore, we observed increased levels of phosphorylated AKT at S473 and phosphorylated ULK1 at S757 (Figure S4F, S4G), indicating increased Akt/mTOR signaling. The increased mTOR activity could be validated by increased levels of pp70S6K (Figure 4F). Finally, loss of FKBP51 resulted in decreased levels of LC3B-II and the accumulation of p62 (Figure 4G, 4H). Together, these data suggest that FKBP51 deletion reduced autophagy signaling in the MBH via the reduction of AMPK activity as well as an increased mTOR signaling, which is in line with our *in vitro* data.

**Fig. 4:**
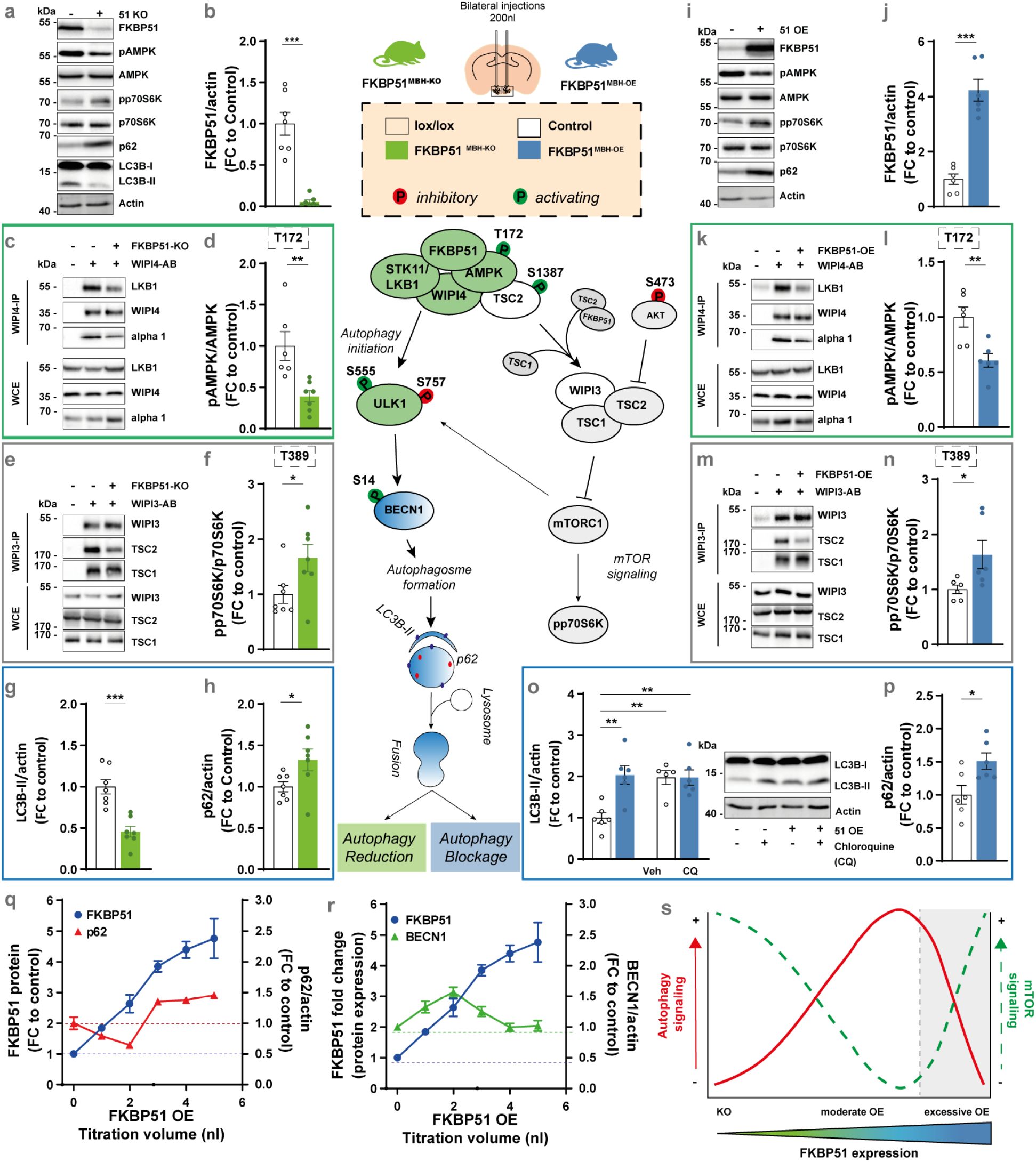
MBH FKBP51 regulates autophagy in an inversed u-shaped manner. FKBP51 deletion is depicted in green and FKBP51 overexpression is depicted in blue. **(A)** Representative blots of autophagy and mTOR markers in FKBP51^MBH-KO^ mice. **(B)** Quantification of FKBP51 deletion. **(C)** FKBP51 deletion reduced LKB1 and AMPK binding to WIPI4 as well as **(D)** AMPK phosphorylation at T172. **(E)** TSC2-WIPI3 binding was decreased in FKBP51^MBH KO^ animals. **(F)** Quantification of mTOR substrate pp70S6K (T389). **(G)** LC3B-II and **(H)** p62 levels in the MBH. **(I)** Representative blots of autophagy and mTOR marker in FKBP51^MBH-OE^ mice. **(J)** Quantification of viral FKBP51 overexpression. **(K)** FKBP51 overexpression reduced LKB1 and AMPK binding to WIPI4. **(L)** Quantification of AMPK phosphorylation at T172. **(M)** TSC2-WIPI3 binding was decreased. **(N)** Quantification of pp70S6K phosphorylation at T389. **(O)** To assess autophagic flux FKBP51^MBH-OE^ animals were treated with 50mg/kg chloroquine and LC3B-II levels were analyzed 4 h after treatment. **(P)** FKBP51 overexpression blocked autophagic flux and resulted in an accumulation of p62. **(Q-R)** Quantification of FKBP51, p62 and BECN1, while titrating AAV-HA-FKBP51 virus into neuroblastoma cells. **(S)** MBH FKBP51 regulates autophagy and mTOR signaling in a dose-dependent manner. All data are shown as ± SEM. Data are shown as the relative protein expression compared to control; for **(A-N)** an unpaired student’s t-test was performed. * p < 0.05, **p < 0.01, ***p < 0.001.

Animals overexpressing FKBP51 in the MBH showed an excessive upregulation of FKBP51 (Figure 4I, 4J). Interestingly, co-IP studies indicated that following the excessive OE of FKBP51, AMPK binding to WIPI4 was decreased. Furthermore, LKB1 levels were strongly reduced and binding of LKB1 to WIPI4 vanished (Figure 4K, Figure S4A). Consequently, phosphorylation of AMPK at T172 was significantly reduced (Figure 4L). Downstream of AMPK, we observed a decrease in phosphorylation of ULK1 at S555 and no changes of phosphorylated BECN1 levels (Figure S4B, S4C). Phosphorylation of TSC2 at S1387 and the binding of TSC2 to WIPI3 was significantly decreased (Figure 4M, Figure S4D-S4E). Phosphorylation of AKT at S473 was unchanged (Figure S4F). On the other hand, FKBP51^MBH-OE^ animals showed increased mTOR signaling, indicated by increased phosphorylation of ULK1 at S757 and elevated levels of pp70S6K (Figure S4G, Figure 4N). Finally, animals overexpressing FKBP51 showed increased levels of LC3B-II. However, treatment with chloroquine (50 mg/kg) did not further increase the LC3B-II levels in FKBP51^MBH-OE^ animals (Figure 4O), indicating that the fusion of autophagosomes with lysosomes is impaired and central autophagic flux is blocked. This hypothesis is supported by the fact that FKBP51 OE resulted in the accumulation of the autophagy substrate p62 (Figure 4P). These data are in contrast to our previously observed findings and imply that FKBP51^MBH-OE^ animals, despite their highly elevated FKBP51 levels, have blocked autophagy signaling in the MBH.

The observation that viral overexpression of FKBP51 in the MBH resulted in a massive OE of FKBP51, led us to hypothesize that the level of FKBP51 expression directly correlates with the degree of autophagy signaling. Therefore, we gradually increased FKBP51 levels in Neuro2a cells by the titration of AAV-HA-FKBP51. Following a moderate increase of FKBP51, we observed a decrease in the accumulation of p62 and an increase in BECN1, supporting the activating role of FKBP51. However, upon a stimulus threshold (at appr. 3-4-fold FKBP51), we observed an inhibitory effect on autophagy with decreased protein level of BECN1 and an increased accumulation of p62 (Figure 4Q, 4R).

Our pathway analysis demonstrates that deletion of FKBP51 reduces autophagy signaling and excessive levels of FKBP51 protein results in a total block of autophagy causing a significant shift from autophagy to mTOR signaling. In conclusion, we suggest that FKBP51 dose-dependently regulates autophagy signaling in an inversed u-shaped manner (Figure 4S).

### MBH FKBP51 alters sympathetic outflow and thereby regulates autophagy signaling in the periphery

The MBH is an established regulatory center for sympathetic outflow to peripheral tissues ^44^. Consequently, we were interested whether the sympathetic tone of the brain into peripheral tissues was affected in FKBP51^MBH-OE^ animals. To do so, we treated FKBP51^MBH-OE^ mice with a single dose of the norepinephrine (NE) synthesis inhibitor α-methyl-p-tyrosine (α-MPT) to block NE synthesis in the periphery, thereby enabling the assessment of the catecholamine turnover rate (cTR) ^45,46^. MBH FKBP51 OE led to a reduction in the cTR in muscle and eWAT (Figure 5A, 5B). Further, inguinal WAT (iWAT) of FKBP51^MBH-OE^ animals showed significant differences in initial NE levels, but not in cTR (Figure S5A). We also observed a mildly but not significantly decreased cTR in brown adipose tissue (BAT) (Figure S5B), whereas no effects were detected in the pancreas or heart tissue (Figure S5C, and S5D). Together, these data demonstrate that MBH FKBP51 OE dampens the sympathetic outflow especially to muscle and fat tissue and encouraged us to investigate changes in autophagy signaling in both tissues.

**Fig. 5:**
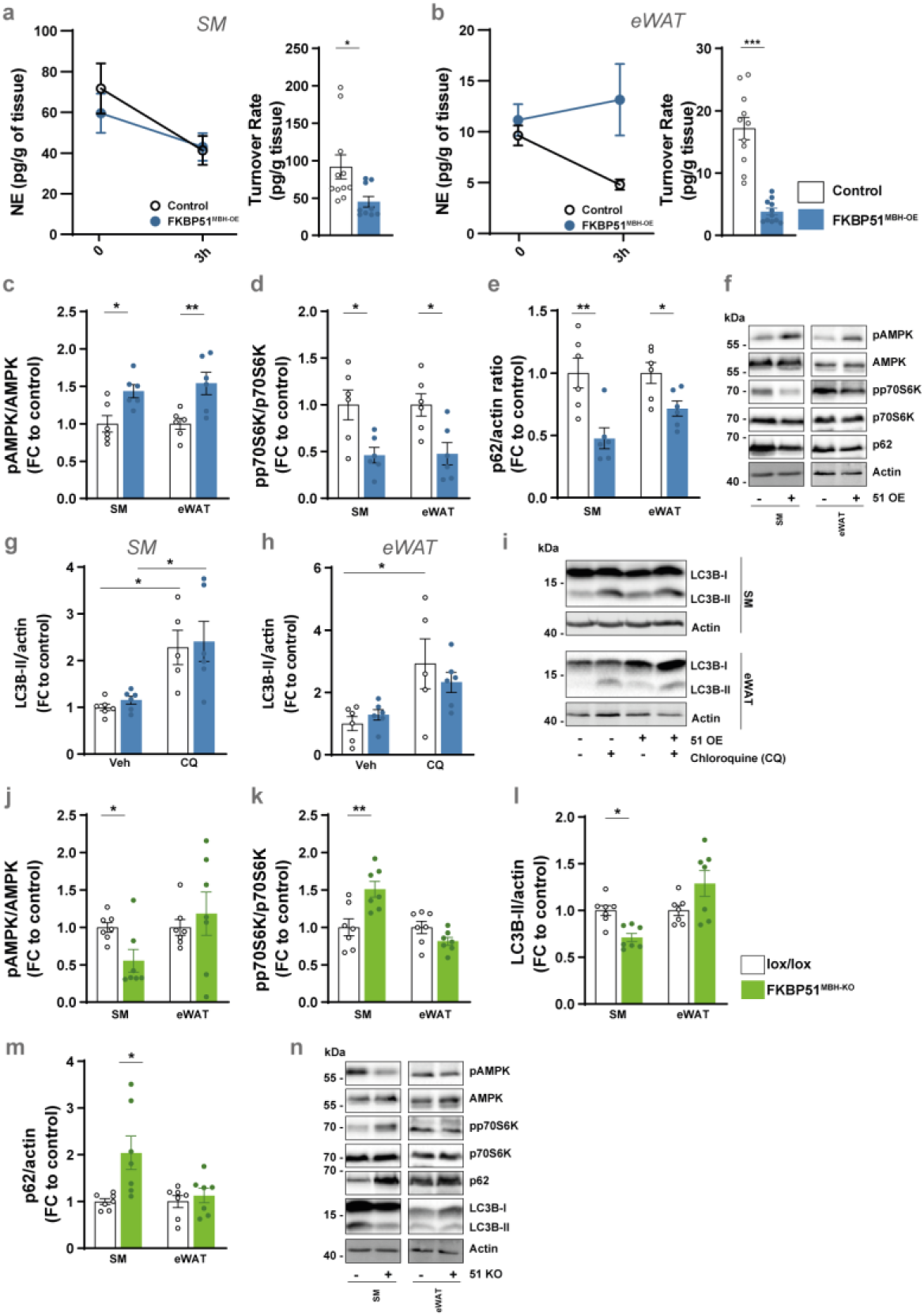
MBH FKBP51 affects sympathetic outflow and peripheral autophagy signaling. FKBP51 overexpression is depicted in blue and FKBP51 deletion is depicted in green. **(A, B)** Representative decrease in tissue NE content after MPT injection (left panel) and turnover rate (right panel) were determined on soleus muscle (SM) and epididymal white adipose tissue (eWAT) (see Figure S3 for pancreas, heart, iWAT and BAT tissues). Quantification of **(C)** pAMPK (T172) and **(D)** pp70S6K (T389), and **(E)** p62 level in the SM and eWAT. **(F)** Representative blots. **(G-H)** FKBP51 overexpression increased autophagic flux and in SM and eWAT. **(I)** Representative blots of chloroquine the experiment. Quantification of **(J)** pAMPK (T172), **(K)** pp70S6K (T389), **(L)** LC3B-II and **(M)** p62 levels in SM and eWAT in animals lacking FKBP51 in the MBH. **(N)** Representative blots of FKBP51^MBH-KO^ protein analysis. All data are shown as ± SEM. Protein data are shown as the relative protein expression compared to control. A two-way ANOVA was performed followed by a Tukey’s multiple comparisons test in **(F-G)**. For **(A - E and I-L)** an unpaired student’s t-test was performed. * p < 0.05, **p < 0.01, ***p < 0.001

In FKBP51^MBH-OE^ animals we observed increased levels of FKBP51 in SM and eWAT (Figure S5E), which resulted in increased AMPK activity (Figure 5C) by enhanced binding of AMPK and LKB1 to WIPI4 (Figure S5F, S5G). Downstream of AMPK, we observed increased phosphorylation of ULK1, BECN1, and TSC2, indicating enhanced autophagy initiation in the periphery (Figure S5H – S5J). We again assessed mTOR signaling and observed increased binding of TSC2 to WIPI3 (Figure S5K) and a strong reduction in the phosphorylation of AKT at S473 and ULK1 at S757 (Figure S5L, S5M), which resulted in reduced levels of pp70S6K (T389) (Figure 5D). Furthermore, we monitored reduced levels of p62 (Figure 5E, 5F). To verify the increase in autophagy signaling, we analyzed LC3B-II levels before and after chloroquine treatment. Here, we detected a true increase in LC3B-II levels after chloroquine treatment (Figure 5G – 5I). These data imply an increase in autophagy flux in the periphery of FKBP51^MBH-OE^ animals. This is in line with our hypothesis that moderately elevated levels of FKBP51 increase autophagy signaling and suggest that the balance between active mTOR signaling in the MBH and active autophagy signaling in the periphery is one driving factor of the lean phenotype of the FKBP51^MBH-OE^ animals.

In FKBP51^MBH-KO^ mice we observed an opposing phenotype with reduced autophagy signaling in SM, whereas autophagy signaling in eWAT was unaltered. In both tissues, we did not observe significant changes in FKBP51 protein level (Figure S5N). However, we could detect less phosphorylation of AMPK at T172 (Figure 5J) and reduced binding of AMPK/LKB1 to WIPI4 (Figure S5O, S5P). These findings were accompanied by reduced levels of ULK1, BECN1, and TSC2 (Figure S5Q-S5S). Furthermore, we monitored increased levels of pp70S6K (T389), pULK1 (S757), and pAKT (S473), suggesting increased mTOR signaling (Figure 5K, Figure S5U, S5V). Finally, LC3B-II levels were significantly reduced (Figure 5L) in combination with elevated levels of p62 in SM (Figure 5M, 5N) which is indicative of reduced autophagy signaling solely in this peripheral tissue. Here, we suggest that the reduction of central and peripheral autophagy signaling is driving the observed body weight phenotype in FKBP51^MBH-KO^ mice.

## Discussion

In the current study, we examined the role of stress-activated chaperone FKBP51 as a molecular master switch linking autophagy and whole-body metabolism. We here present that FKBP51 actively modulates the response of the AMPK-mTOR network to a high-fat diet by scaffolding autophagy-upstream AMPK/LKB1/WIPI4 and TSC2/WIPI3 hetero-protein complexes. Importantly, we identify a tissue-specific function of FKBP51 by providing *in vivo* evidence that hypothalamic FKBP51 acts as a dose-specific mediator of whole-body metabolism.

Metabolomic profiling of neuronal-like cells lacking FKBP51 revealed a substantial increase for several metabolites and amino acids and suggests a role of FKBP51 in BCAA metabolism. BCAAs are important regulators of neurotransmitters and protein synthesis as well as food intake ^47,48^. The increase of multiple BCAAs have been associated with obesity and insulin resistance ^49,50^. In our *in vitro* metabolomic profiling analysis, isoleucine, leucine, valine, and tyrosine were strongly elevated in FKBP51 KO cells under normal and high glucose concentrations, which is indicative of constantly active mTOR signaling ^51^. Furthermore, it has been shown that excess leucine can reduce abdominal fat loss whereas leucine deprivation promotes fat loss via CREB (cAMP response element-binding protein) signaling and increased expression of CRH (corticotropin-releasing hormone) in the hypothalamus. This effect is conveyed by the activation of the sympathetic nervous system ^52^. Leucine is an important regulator of mTOR and negatively affects the biogenesis of autophagosomes through its metabolite acetyl CoA, which thereby enhances acetylation of the mTOR associated protein RPTOR via acetyltransferase EP300 in neurons and other cell types. This cascade of events ultimately leads to autophagy inhibition and mTOR activation ^53,54^. At the same time, the increased levels of polyamines, observed in FKBP51 KO cells, are in contrast to autophagy inhibition. Especially spermidine was shown to be capable of autophagy induction via inhibition of EP300 ^55^. Nevertheless, cell-type specific effects have to be taken into account as studies suggest an increased expression of EP300 in response to spermidine supplementation in aged and osteoarthritic chondrocytes ^56^.

To gain further insight into the underlying mechanisms, we built on already existing knowledge about FKBP51 regarding its regulatory role on single autophagy-related proteins (like BECN1, WIPIs, and SKP2 ^11,16,26^) which further positions FKBP51 as a major upstream regulator of autophagy. AMPK is activated by the phosphorylation of T172, which is regulated by LKB1 ^57^ and increased LKB1/AMPK signaling activates the TSC1/TSC2 complex, which in turn inhibits mTOR activity ^40^. Recently, Bakula and colleagues showed that the WIPI protein family members WIPI3-WIPI4 are essential scaffolders of the LKB1/AMPK/TSC1/2 signaling network thereby regulating autophagy and mTOR signaling ^11^. Here, we extended this knowledge by revealing that FKBP51 recruits LKB1 to the WIPI4-AMPK regulatory platform to induce AMPK phosphorylation at T172, which further increases autophagy initiation by direct phosphorylation of ULK1 at S555 ^10^. On the other hand, FKBP51 associates with the TSC2/WIPI3 heterocomplex to co-regulate mTOR signaling and thus position FKBP51 as a main regulatory switch between autophagy initiation and mTOR signaling. In fact, our experiments *in vivo* and *in vitro* led us to propose a model in which physiological levels of FKBP51 are essential for normal cellular autophagy-mediated homeostasis. Hereby, the absence of FKBP51 reduces autophagy signaling and capacity in contrast to excessive, non-physiological levels of FKBP51, which block autophagy in favor of mTOR signaling. The relative amounts of FKBP51 complexed with autophagy regulators within a cell govern the threshold for a transition from cell homeostasis to impaired autophagy in an inversed u-shaped manner.

FKBP51 has been shown to act tissue-specific to control adipocyte differentiation, browning ^19,58^, and glucose metabolism ^18^. Most studies investigated global FKBP51 KO mice and observed a lean phenotype after a HFD regimen. These observations would imply that hypothalamic FKBP51 OE increases body weight gain whereas deletion reduces body weight. Remarkably, we here observed that the acute FKBP51 manipulation in the MBH acts in an opposing manner with a central deletion leading to obesity and an overexpression to a lean phenotype, emphasizing its tissue specificity. These data suggest that hypothalamic FKBP51 regulates body weight and food intake in a u-shaped manner, which is supported by the fact that HFD increases FKBP51 expression and a modest increase of hypothalamic FKBP51 induces body weight gain ^30^.

Several studies have already addressed the functional role of hypothalamic autophagy in the regulation of whole-body metabolism. Mice challenged with a chronic HFD showed impaired autophagy in the arcuate nucleus (ARC) and a deletion of ATG7 in the MBH resulted in hyperphagia and increased body weight ^59^. The MBH, however, is a complex brain region with multiple different nuclei, which have various opposing roles in the control of whole-body metabolism ^60,61^. Here, we targeted FKBP51 non-selectively in various nuclei within the MBH, which is a limitation of the study. However, it also raises the question, which nuclei or neuronal subpopulations are the driving force behind our observed phenotypes. Previous studies have shown that specific deletion of autophagy in POMC neurons leads to hyperphagia and obesity ^12^. These findings are in line with our data of the FKBP51^MBH-KO^ animals, which develop obesity and hyperphagia on a regular chow diet in combination with decreased activity of central autophagy. Intriguingly, animals overexpressing FKBP51 in the MBH have a reduced body weight progression, increased glucose tolerance, and insulin sensitivity under a HFD regimen, despite the hypothalamic autophagy blockage. Interestingly, the deletion of autophagy in AgRP neurons resulted in decreased body weight and food intake in response to fasting ^13^. Our lab’s unpublished observations suggest a high and distinct expression of FKBP51 within specific neuronal subpopulations of the ARC (unpublished data), which may suggest an overlapping role of FKBP51 and autophagy here. Future studies should emphasize the specific neuronal action of FKBP51 on autophagy to regulate body weight progression and food intake.

Finally, we suggest that the altered balance between hypothalamic and peripheral autophagy-mTOR signaling is a major contributor to the observed phenotype of FKBP51^MBH-OE^ and FKBP51^MBH-KO^ animals. Indeed, FKBP51^MBH-KO^ animals showed decreased autophagy and increased mTOR signaling in the periphery (eWAT and SM), whereas peripheral autophagy signaling in FKBP51^MBH-OE^ animals displayed the opposite phenotype. The importance of central-peripheral mTOR and autophagy signaling has already been studied intensely. For instance, peripheral mTOR activity was shown to be involved in the pathogenesis of obesity and is enhanced in muscle and adipose tissue in obese animals ^62,63^, whereas the activation of central mTOR can reduce food intake and body weight gain ^64^. In particular, the mTOR substrate p70S6K was shown to regulate body weight by mediating the sensitivity of leptin to AMPK via PI3K/AKT1/mTOR pathway ^65,66^. Here, we extend this finding by the fact that the combination of peripheral and central p70S6K activity is an important contributor to the development of obesity.

In conclusion, this study provides a new conceptual framework for the regulatory function of the stress responsive co-chaperone FKBP51 on autophagy signaling and establishes a physiological role of MBH FKBP51 in the regulation of food intake and body weight regulation. We further suggest that FKBP51 is a crucial sensor linking signaling pathways controlling the stress response, autophagy, and metabolism. The newly discovered ability of FKBP51 to regulate autophagy and energy homeostasis might therefore open new promising treatment avenues for metabolic disorders, such as obesity and type 2 diabetes.

## Methods

### Antibodies

Goat polyclonal anti-actin, (I-19)(sc-1616, Santa Cruz Biotechnology), Rabbit polyclonal anti-FKBP51 (A301-430A, Bethyl Laboratories), Rabbit monoclonal anti-LKB1 (D60C5, #3047, Cell Signaling Technology), Rabbit polyclonal anti-pAMPKα^T172^ (#2531, Cell Signaling Technology), Rabbit polyclonal anti-pAMPKα (#2532, Cell Signaling Technology), Rabbit polyclonal anti-SKP2, L70, (#4313, Cell Signaling Technology), Rabbit anti-pSKP2^S72^ (was a kind gift from Cell Signaling Technology), Rabbit polyclonal anti-AKT (#9272, Cell Signaling Technology), Rabbit monoclonal anti-pAKT^S473^ (D9E, #4060, Cell Signaling Technology), Rabbit polyclonal anti-p62 (#5114, Cell Signaling Technology), Rabbit monoclonal anti-LC3B (D11, #3868, Cell Signaling Technology), Rabbit polyclonal anti-pULK1^S757^ (#6888, Cell Signaling Technology), Rabbit monoclonal anti-pULK1^S555^ (D1H4, #5869, Cell Signaling Technology), Rabbit monoclonal anti-ULK1 (D8H5, #8054, Cell Signaling Technology), anti-pBECN1^S93/S96^, (in mouse S91/S94) (#12476, Cell Signaling Technology), Rabbit polyclonal anti-pBECN1^S15^ (#84966, Cell Signaling Technology), Rabbit polyclonal anti-BECN1 (#3738, Cell Signaling Technology), Rabbit polyclonal anti-TSC2 (#3612, Cell Signaling Technology), Rabbit polyclonal anti-pTSC2^S1387^ (#5584, Cell Signaling Technology), Rabbit monoclonal anti-pATG16L1^S278^ (EPR19016, ab195242, abcam), Rabbit polyclonal anti-WIPI4 (WDR45) (19194-1-AP, ProteinTech), Rabbit polyclonal anti-WIPI3 (WDR45L) (SAB2102704, Sigma), Rabbit polyclonal anti-WIPI2 (#8567, Cell Signaling Technology), Rabbit polyclonal anti-WIPI1 (HPA007493, Sigma), Rabbit polyclonal anti-AMPKα1 (#2795, Cell Signaling Technology), Rabbit polyclonal anti-AMPKγ2 (#2536, Cell Signaling Technology), Rabbit polyclonal anti-AMPKα2 (#2757, Cell Signaling Technology), Rabbit monoclonal anti-AMPKβ1 (71C10, #4178, Cell Signaling Technology), Rabbit polyclonal anti-AMPKγ1 (#4187, Cell Signaling Technology), Rabbit polyclonal anti-AMPKβ2 (#4188, Cell Signaling Technology), Rabbit polyclonal anti-AMPKγ3 (#2550, Cell Signaling Technology), Rabbit monoclonal anti-TSC1 (D43E2, #6935, Cell Signaling Technology), Rabbit polyclonal anti-FLAG (600-401-383, Rockland Inc.).

### Animals & animal housing

All experiments and protocols were approved by the committee for the Care and Use of Laboratory animals of the Government of Upper Bavaria and were performed in accordance with the European Communities’ Council Directive 2010/63/EU. All animals were kept singly housed in individually ventilated cages (IVC; 30cm × 16 cm × 16 cm; 501 cm^2^) with *ad libitum* access to water and food and constant environmental conditions (12:12 hr light/dark cycle, 23±2°C and humidity of 55%) during all times. All IVCs had sufficient bedding and nesting material as well as a wooden tunnel for environmental enrichment. All animals were fed with a standard research chow diet (Altromin 1318, Altromin GmbH, Germany) or a high fat diet (HFD, 58% kcal from fat, D12331, Research Diets, New Brunswick, NJ, USA). For all experiments, male C57Bl/6n or male *Fkbp5*^lox/lox^ mice (described in Ref ^67^) aged between 2-5 months were used.

### Viral overexpression and knockdown of *FKBP51*

For overexpression of FKBP51, we injected an AAV vector containing a CAG-HA-tagged-FKBP51-WPRE-BGH-polyA expression cassette (containing the coding sequence of human FKBP51 NCBI CCDS ID CCDS4808.1) in C57Bl/6n mice. The same vector construct without expression of Fkbp5 (CAG-Null/Empty-WPRE-BGH-polyA) was used as a control. Virus production, amplification, and purification were performed by GeneDetect. A viral vector containing a *Cre* expressing cassette (pAAV-CMV-HI-eGFP-Cre-WPRE-SV40, Addgene; #105545) was used to induce *Fkbp5* deletion in *Fkbp5*^lox/lox^ mice. Control animals were injected with a control virus (pAAV-CMV-PI-eGFP-WPRE-bGH; Addgene; #105530). For both experiments, stereotactic injections were performed as described previously ^68^. In brief, mice were anesthetized with isoflurane prior surgery, and 0.2 μl of the above-mentioned viruses (titers: 1.6 × 10^12-13^ genomic particles/ml) were bilaterally injected in the MBH at 0.05 μl/min by glass capillaries with tip resistance of 2–4MΩ in a stereotactic apparatus. The following coordinates were used: −1.5 mm anterior to bregma, 0.4 mm lateral from midline, and 5.6 mm below the surface of the skull, targeting the MBH. After surgery, mice were treated for 3 d with Metacam via i.p. injections and were housed for 3-4 weeks for total recovery prior the actual experiments. Successful overexpression and knockdown of *Fkbp5* was verified by in-situ hybridization and western blot.

### Autophagic flux

We investigated the autophagic flux by the injection of chloroquine (50 mg/kg,), an inhibitor of lysosomal acidification and autophagosome-lysosomal fusion that blocks degradation of autophagosome cargo ^37^. We injected C57Bl/6n mice in the morning with 50 mg/kg chloroquine or saline as control. Multiple tissues were removed and shock frozen 4 hours after injection and stored at −80°C until protein analysis.

### Sample collection

On the day of sacrifice, animals were deeply anesthetized with isoflurane and sacrificed by decapitation. Trunk blood was collected in labeled 1.5 ml EDTA-coated microcentrifuge tubes (Sarstedt, Germany) and kept on ice until centrifugation. After centrifugation (4°C, 8000rpm for 1 min) the plasma was removed and transferred to new, labeled tubes and stored at −20°C until hormone quantification. For mRNA analysis, brains were removed, snap-frozen in isopentane at −40°C and stored at −80°C for ISH. For protein analysis, the mediobasal hypothalamus, skeletal muscle (soleus muscle), and WAT (eWAT) were collected and immediately shock frozen and stored at −80°C until protein analysis.

### Catecholamine turnover rate determination

Catecholamine turnover was measured on the basis of the decline in tissue norepinephrine (NE) content after the inhibition of catecholamine biosynthesis with α-methyl-DL-tyrosine methyl ester hydrochloride ((α-MPT, Sigma Aldrich, ST, Quentin, France) 200 mg/kg i.p. injection), as described previously ^45^.

In the morning, bedding was changed, and C57Bl/6n mice were food deprived for 3h to insure post-prandial state and injected with α-methyl-DL-tyrosine (MPT; a tyrosine hydroxylase inhibitor) to block catecholamine synthesis. Prior to (time = 0) and 3h after the injection (time = 3h), animals were sacrificed, the tissues were removed, flash frozen in liquid nitrogen and stored at −80°C for monoamine and metabolite analysis.

Catecholamine content at time = 0 (NE (0)) was determined on a group of animals receiving a saline injection. Because the concentration of catecholamine in tissues declined exponentially, we could obtain the rate constant of NE efflux (expressed in h-1). Comprehensive analysis of norepinephrine was carried out by reverse-phase liquid chromatography with electrochemical detection as described by ^69^. The values obtained were expressed as ng/mg wet tissue and were logarithmically transformed for calculation of linearity of regression, standard error of the regression coefficients and significance of differences between regression coefficients.

### Glucose tolerance and insulin tolerance

Alteration of glucose metabolism in FKBP51^MBH OE^ and FKBP51^MBH KO^ mice were investigated by a glucose (GTT) and insulin tolerance (ITT) test as described previously ^18^.

### Hormone assessment

Corticosterone concentrations were determined by radioimmunoassay using a corticosterone double antibody ^125^I RIA kit (sensitivity: 12.5 ng/ml, MP Biomedicals Inc) and were used following the manufacturers’ instructions. Radioactivity of the pellet was measured with a gamma counter (Packard Cobra II Auto Gamma; Perkin-Elmer). Final CORT levels were derived from the standard curve.

### Cell lines

Neuro2a WT, SH-SY5Y WT and FKBP51KO ^70^ cells were maintained in Dulbecco’s Modified Eagle’s Medium (DMEM) supplemented with 10% fetal bovine serum and 1x penicillin-streptomycin antibiotics at 37 °C in a humidified atmosphere with 5% CO_2_. At 90% confluency, Neuro2a cells were detached from the plate and 2×10^6^ cells were resuspended in 100 μl of transfection buffer (50 nM HEPES [pH 7.3], 90 mM NaCl, 5 mM KCl, and 0.15 mM CaCl2). A total of 2.5 μg of plasmid DNA or 80 ng of siRNA (siWIPI3, EMU081491 or siWIPI4, EMU007321 or siControl, SIC001 all Sigma) was used per transfection. Electroporation was performed using the Amaxa Nucleofector system 2b (program #T-020).

### Co-immunoprecipitation (coIP)

Immunoprecipitations of endogenous proteins were performed from protein extracts (n = 3-4 per group) derived from Neuro2a cells, SH-SY5Y WT or FKBP51KO cells, soleus muscle, eWAT and MBH. For Co-IPs, 500 μg of lysate was incubated with 2 μg of the appropriate IP antibody (anti-FLAG (FKBP51), anti-WIPI4, anti-WIPI3) at 4 °C overnight. 20 μl of rabbit IgG-conjugated protein G dynabeads (Invitrogen, 100-03D) were blocked with BSA and subsequently added to the lysate-antibody mixture and allowed to incubate at 4 °C for 3 h in order mediate binding between dynabeads and the antibody-antigen complex of interest. Beads were then washed three times with ice-cold PBS and the protein antibody complexes eluted with 60 μl Laemmli loading buffer. Thereafter, the eluate was boiled for 5 min at 95 °C. Then, 2 to 5 μl of each immunoprecipitate was separated by SDS-PAGE and electro-transferred onto nitrocellulose membranes. For assessing protein complexes, immunoblotting against WIPI1-4, FKBP51, LKB1, AMPK, TSC1 and TSC2 was performed.

### Western blot analysis

Protein extracts were obtained by lysing cells (in RIPA buffer (150 mM NaCl, 1% IGEPAL CA-630, 0.5% Sodium deoxycholate, 0.1% SDS 50mM Tris (pH8.0)) freshly supplemented with protease inhibitor (Merck Millipore, Darmstadt, Germany), benzonase (Merck Millipore), 5 mM DTT (Sigma Aldrich, Munich, Germany), and phosphatase inhibitor cocktail (Roche, Penzberg, Germany). Proteins were separated by SDS-PAGE and electro-transferred onto nitrocellulose membranes. Blots were placed in Tris-buffered saline, supplemented with 0.05% Tween (Sigma Aldrich) and 5% non-fat milk for 1 h at room temperature and then incubated with primary antibody (diluted in TBS/0.05% Tween) overnight at 4°C.

Subsequently, blots were washed and probed with the respective horseradish peroxidase or fluorophore-conjugated secondary antibody for 1 h at room temperature. The immuno-reactive bands were visualized either using ECL detection reagent (Millipore, Billerica, MA, USA) or directly by excitation of the respective fluorophore. Determination of the band intensities were performed with BioRad, ChemiDoc MP.

### LC-MS analysis of amine-containing metabolites

**T**he benzoyl chloride derivatization method was used for amino acid analysis ^71^. In brief: the dried metabolite pellets were resuspended in 90 μl of the LC-MS-grade water (Milli-Q 7000 equipped with an LC-Pak and a Millipak filter, Millipore). Then, 20 μl of the resuspended sample was mixed with 10 μl of 100 mM sodium carbonate (Sigma-Aldrich) followed by the addition of 10 μl 2% benzoyl chloride (Sigma-Aldrich) in acetonitrile (Optima-Grade, Fisher-Scientific). Samples were vortexed before centrifugation for 10 min at 21,300 × g at 20°C. Clear supernatants were diluted 1:10 with LC MS-grade water and transferred to fresh auto-sampler tubes with conical glass inserts (Chromatographie Zubehoer Trott) and analyzed using a Vanquish UHPLC (Thermo) connected to a Q-Exactive HF (Thermo).

For the analysis, 1 μl of the derivatized sample were injected onto a 100 × 2.1 mm HSS T3 UPLC column (Waters). The flow rate was set to 400 μL/min using a buffer system consisting of buffer A (10 mM ammonium formate (Sigma-Aldrich), 0.15% formic acid (Sigma-Aldrich) in LC MS-grade water), and buffer B (acetonitrile, Optima-grade, Fisher-Scientific). The LC gradient was: 0% B at 0 min; 0-15% B 0-0.1 min; 15-17% B 0.1-0.5 min; 17-55% B 0.5-7 min, 55-70% B 7-7.5 min; 70-100% B 7.5-9 min; 100% B 9-10 min; 100-0% B 10-10.1 min, 10.1-15 min 0% B. The mass spectrometer was operating in positive ionization mode monitoring the mass range m/z 50-750. The heated ESI source settings of the mass spectrometer were: Spray voltage 3.5 kV, capillary temperature 250°C, sheath gas flow 60 AU and aux gas flow 20 AU at a temperature of 250°C. The S-lens was set to a value of 60 AU.

Data analysis was performed using the TraceFinder software (Version 4.2, Thermo Fisher Scientific). Identity of each compound was validated by authentic reference compounds, which were analyzed independently. Peak areas were analyzed by using extracted ion chromatograms (XIC) of compound-specific [M + nBz + H]^+^ where n corresponds to the number of amine moieties, which can be derivatized with a benzoyl chloride (Bz). XIC peaks were extracted with a mass accuracy (<5 ppm) and a retention time (RT) tolerance of 0.2 min.

### Anion-Exchange Chromatography Mass Spectrometry (AEX-MS) of the analysis of TCA cycle and glycolysis metabolites

Anion-Exchange Chromatography was performed simultaneously to the LC MS analysis.First, 50 μl of the re-suspended sample were diluted 1:5 with LC MS-grade water and analyzed using a Dionex ion chromatography system (ICS 5000, Thermo Scientific). The applied protocol was adopted from ^72^. In brief: 10 μL of polar metabolite extract were injected in full loop mode using an overfill factor of 3, onto a Dionex IonPac AS11-HC column (2 mm × 250 mm, 4 μm particle size, Thermo Scientific) equipped with a Dionex IonPac AS11-HC guard column (2 mm × 50 mm, 4 μm, Thermo Scientific). The column temperature was held at 30°C, while the auto-sampler was set to 6°C. A potassium hydroxide gradient was generated by the eluent generator using a potassium hydroxide cartridge that was supplied with deionized water. The metabolite separation was carried at a flow rate of 380 μL/min, applying the following gradient: 0-5 min, 10−25 mM KOH; 5-21 min, 25−35 mM KOH; 21-25 min, 35-100 mM KOH, 25-28 min, 100 mM KOH, 28-32 min, 100-10 mM KOH. The column was re-equilibrated at 10 mM for 6 min. The eluting metabolites were detected in negative ion mode using ESI MRM (multi reaction monitoring) on a Xevo TQ (Waters) triple quadrupole mass spectrometer applying the following settings: capillary voltage 1.5 kV, desolvation temperature 550°C, desolvation gas flow 800 L/h, collision cell gas flow 0.15 mL/min. All peaks were validated using two MRM transitions, one for quantification of the compound, while the second ion was used for qualification of the identity of the compound. Data analysis and peak integration was performed using the TargetLynx Software (Waters).

### Statistical analysis

The data presented are shown as means ±SEM and samples sizes are indicated in the figure legends. All data were analyzed by the commercially available software SPSS v17.0 and GraphPad v8.0. The unpaired student’s t-test was used when two groups were compared. For four group comparisons, two-way analysis of variance (ANOVA) was performed, followed by Tukey’s multiple comparisons test, as appropriate. P values of less than 0.05 were considered statistically significant.

### Data availability

The data herein are available from the corresponding authors upon reasonable request.

## Acknowledgments

The authors thank Claudia Kühne, Daniela Harbich and Bianca Schmid (Max Planck Institute of Psychiatry, Munich, Germany) for their excellent technical assistant and support. We thank Jan Deussing, and the scientific core unit *Genetically Engineered Mouse Models* for providing technical support and guidance. Funding: This work was supported by the “OptiMD” grant of the Federal Ministry of Education and Research (01EE1401D; M.V.S.), the BioM M4 award “PROCERA” of the Bavarian State Ministry (M.V.S.), the “GUTMOM” grant of the Federal Ministry of Education and Research (01EA1805; M.V.S.). and the “Kids2Health” grant of the Federal Ministry of Education and Research (01GL1743C; M.V.S.).

## Author contributions

A.S.H, G.B., M.V.S., and N.C.G.: Conceived the project and designed the experiments. A.S.H and L.M.B. managed the mouse lines and genotyping. A.S.H., M.L.P., and L.M.B. performed animal experiments and surgeries. K.H., T.B., and N.C.G.: Performed protein work. P.G. and T.B.: Performed and analyzed metabolomics experiments. M.D.A, J.N., and KW. S.: Performed catecholamine extraction. A.S.H.: Wrote the initial version of the manuscript. M.V.S., A.C, and N.C.G.: Supervised the research and all authors revised the manuscript.

## Supplementary Information

**Fig. S1:**
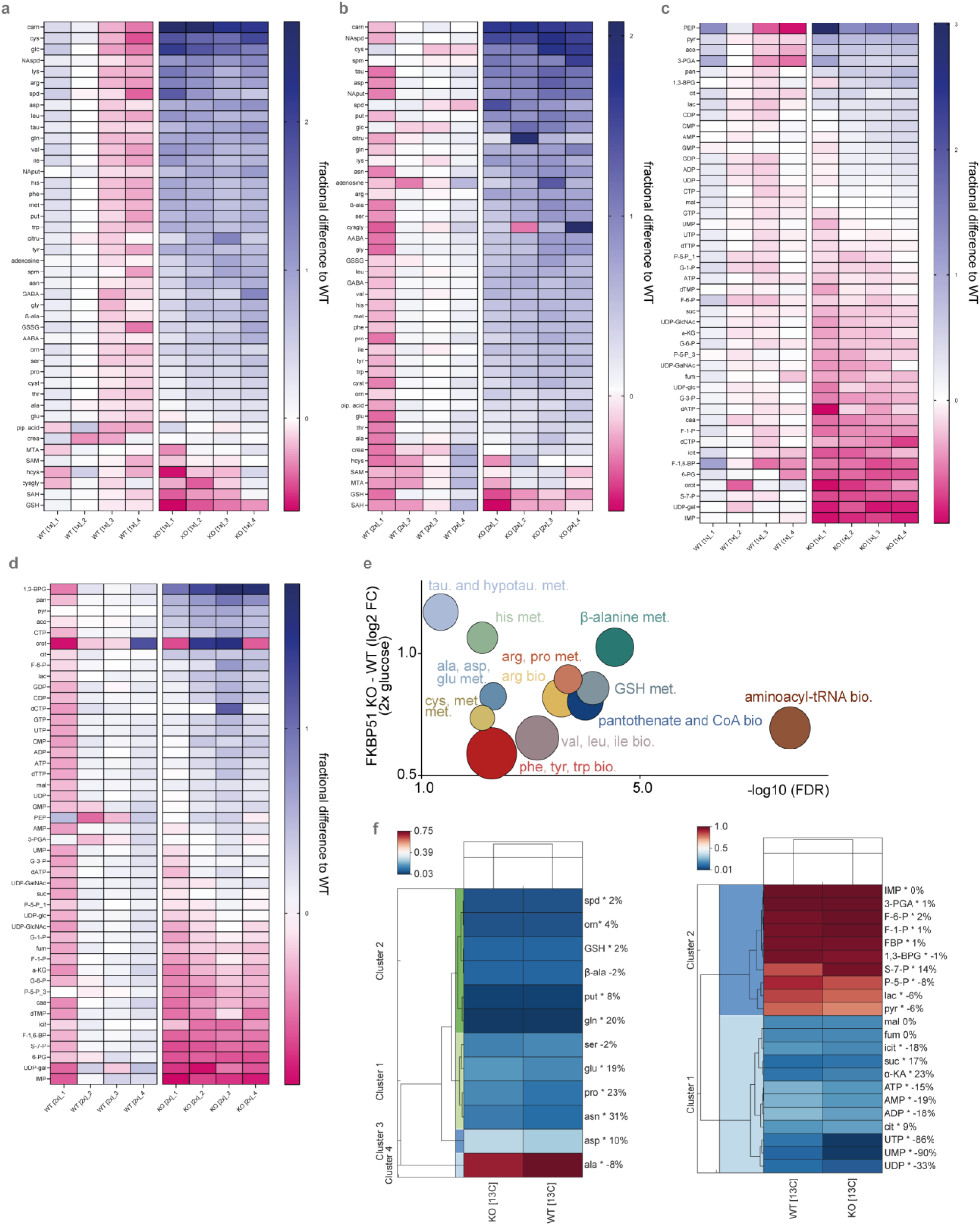
FKBP51 deletion alters AMPK and mTOR associated amino acid metabolic and biosynthetic pathways. **A)** Heatmap of altered amine-containing (Bz) metabolites in SH-SY5Y cells lacking FKBP51 and WT control cells cultured under normal glucose condition (1x, 4.5g/l) and **(B)** increased glucose condition (2x, 9g/l). **(C)** Heatmaps of altered anionic (IC) metabolites in SH-SY5Y cells lacking FKBP51 compared to WT control cells under normal and **(D)** increased glucose culturing conditions. The fractional differences of each replicate are shown for all metabolites comparing the genotype and the different glucose conditions. **(E)** Analysis and regulation of significantly altered pathways of FKBP51 KO and WT cells under excessive glucose conditions. The f(x)-axis shows the (median) log2 fold change (FC) of all significantly altered metabolites of the indicated pathway and the false discovery rate (FDR, equals the –log10 adjusted p-value) is shown on the x-axis. The size of the circles represents the amount of significantly changed metabolites in comparison to all metabolites of a particular pathway. **(F)** Analysis of the metabolic flux in FKBP51 KO cells compared to WT cells using C13 glucose as tracer. The enrichment of C13 is displayed for each metabolite investigated.

**Fig. S2:**
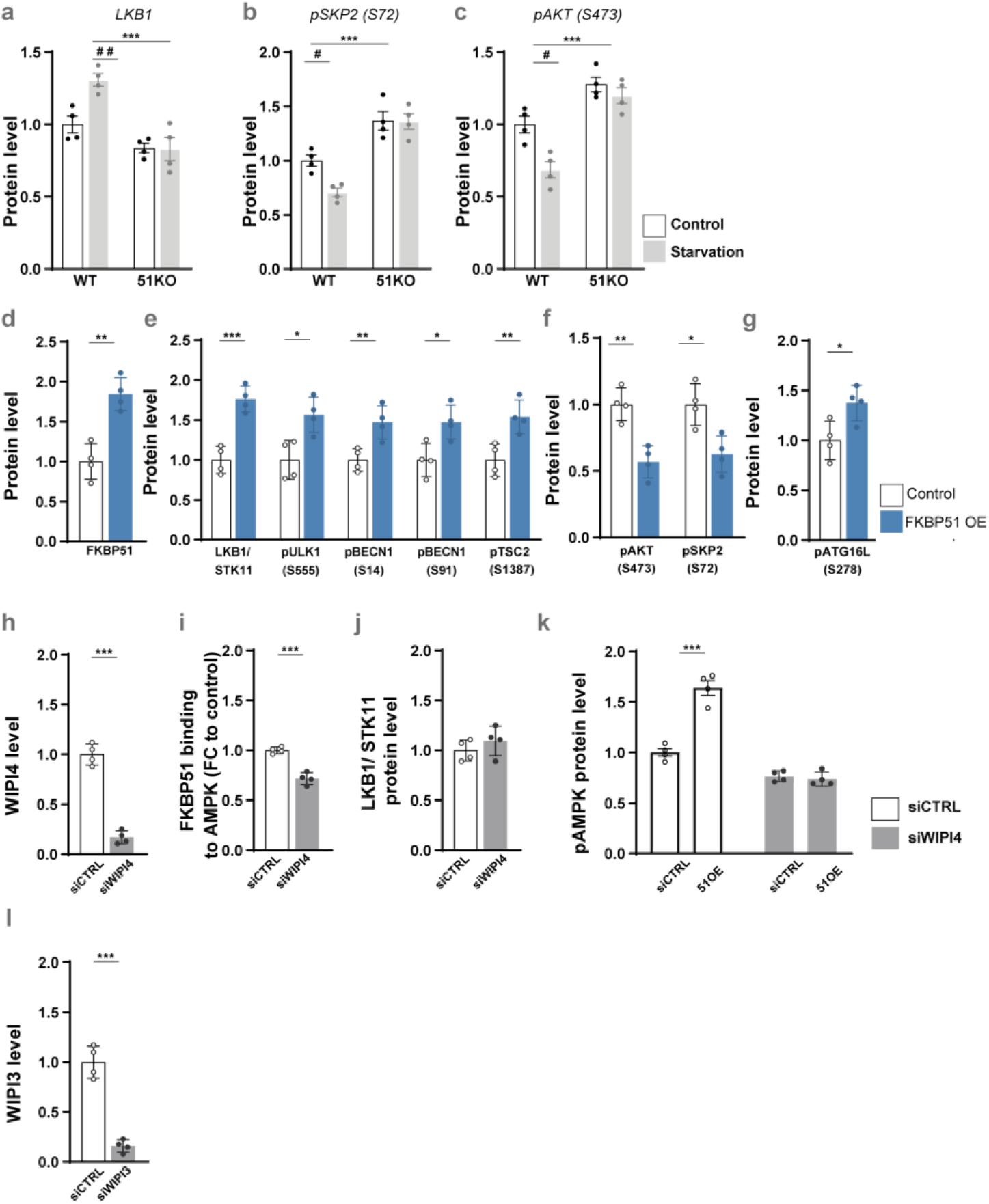
In-vitro manipulation of FKBP51 and its effects on autophagy signaling. Wildtype (WT) or FKBP51 knockout (51KO) cells were starved in HBSS medium for 4 h to induce autophagy. The levels of autophagy markers were determined by immunoblotting. **(A)** Quantification of LKB1, **(B)** pSkp2 (S72), and **(C)** pAkt (S473). **(D)** Validation of FKBP51 overexpression (FKBP51 OE) in neuroblastoma cells**. (E)** FKBP51 OE in Neuro2a cells enhanced phosphorylation of autophagy markers regulating autophagy initiation. **(F)** Quantification of pAKT (S473) and pSKP2 (S72). **(G)** Phosphorylation of ATGL16L at S278. **(H)** Confirmation of WIPI4 knockdown in Neuro2cells. **(I)** FKBP51 binding to AMPKa1 in WIPI4 knockout cells. **(J)** LKB1 binding to FKBP51 were not affected in WIPI4 lacking cells**. (K)** WIPI4 deletion blocked the FKBP51 overexpressing effect on pAMPK at T172. **(L)** WIPI3 knockdown in Neuro2a cells. All data are shown as relative fold change compared to control condition; ± s.e.m.; a two-way ANOVA was perfomed in **(A-C)** and followed by a Tukey’s multiple comparison test. The unpaired student’s t-test was performed in **(D-L)**. * p < 0.05, **p < 0.01, ***p < 0.001; # p < 0.05, ## p < 0.01. * = significant treatment effect; # = significant genotype effect.

**Fig. S3:**
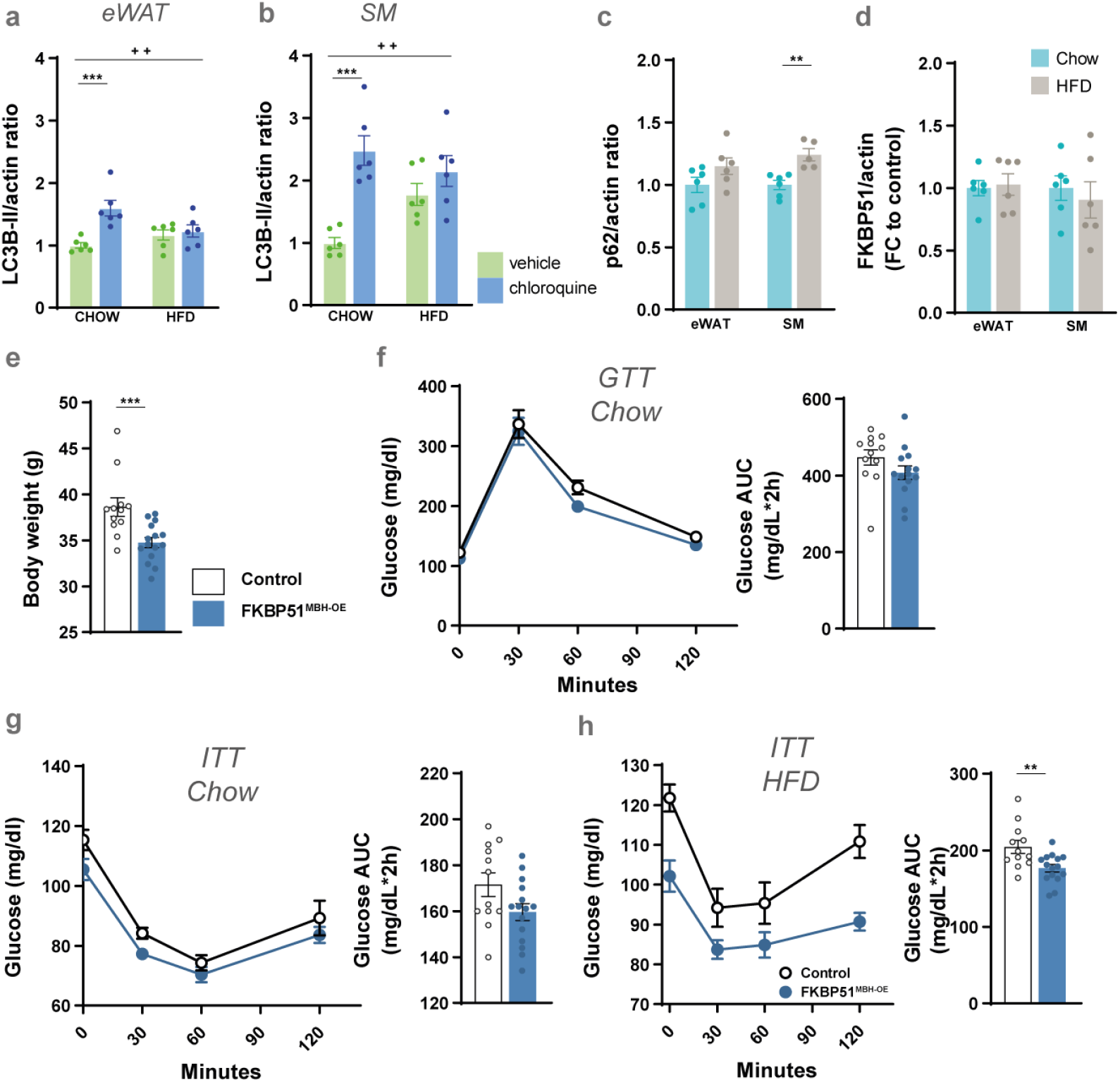
FKBP51 overexpression in the MBH affects sympathetic outflow to muscle and fat tissue. **(A)** LC3B-II level before and after chloroquine treatment (50mg/kg) in eWAT and **(B)** soleus muscle under chow and HFD conditions **(C)**10 weeks of HFD increased the accumulation of the autophagy receptor p62 in SM, but not in eWAT. **(D)** FKBP51 expression in soleus muscle (SM) and epididymal white adipose tissue (eWAT) after 10 weeks of HFD. **(E)** Overexpression of FKBP51 in a second cohort of C57/Bl6 animals resulted in a lean body weight phenotype after 10 weeks of HFD**. (F-H)** Differences in glucose metabolism were investigated by performing a glucose tolerance test (GTT) and an insulin tolerance test (ITT) under chow and HFD conditions. For **(A, B)** a two-way ANOVA was performed followed by a Tukey’s multiple comparisons test and data are shown as relative fold change compared to control condition. For **(C - H)** an unpaired student’s t-test was performed. ± SEM; * p < 0.05, **p < 0.01, ***p < 0.001; # p < 0.05, ## p < 0.01. * = significant treatment effect; + = significant treatment × diet interaction.

**Fig. S4:**
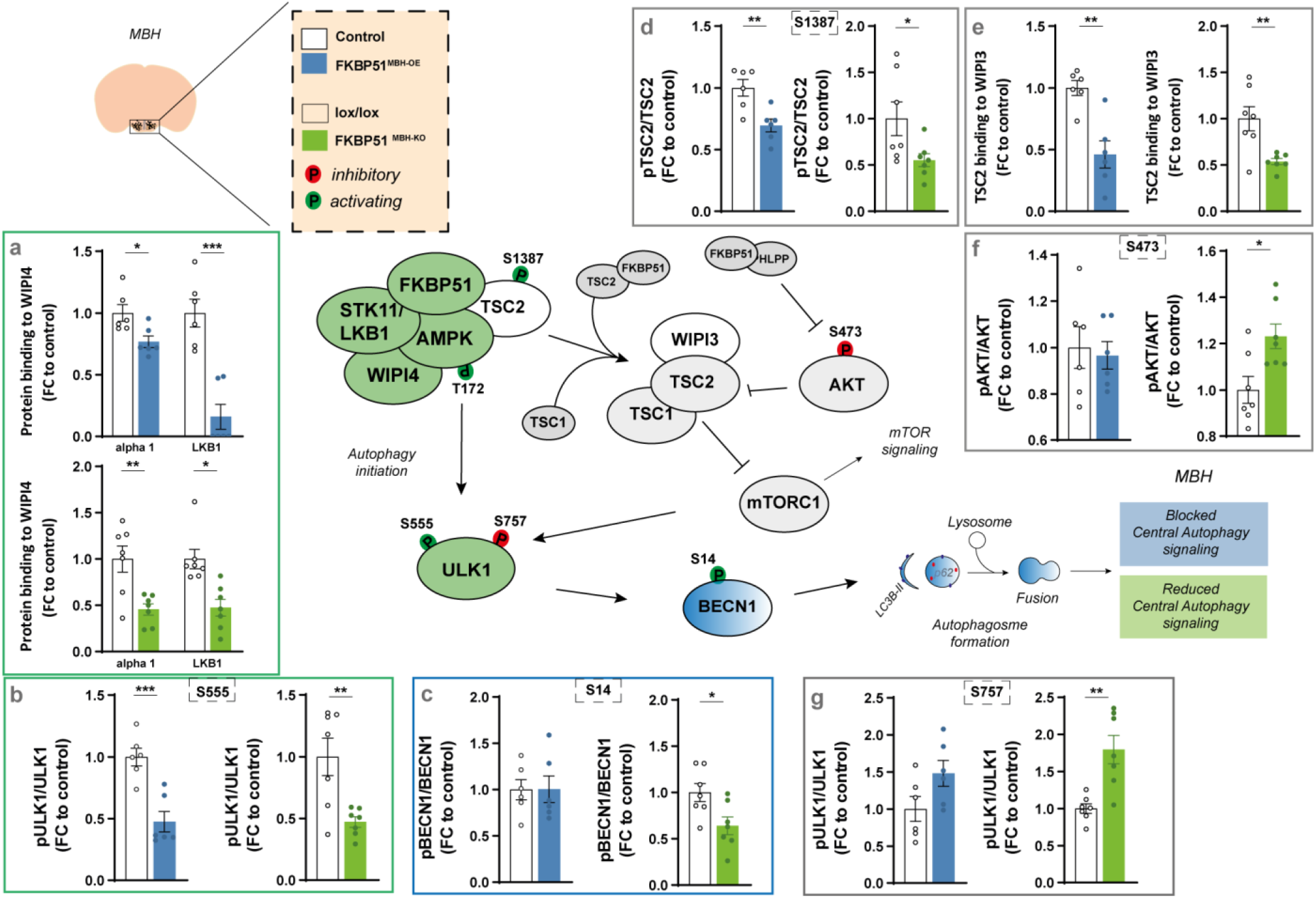
FKBP51 regulates autophagy signaling in the MBH. Pathway analysis of main autophagy and mTOR marker in the mediobasal hypothalamis (MBH). FKBP51 overexpression is depicted in blue and FKBP51 deletion is depicted in green. **(A)** Quantification of LKB1 and AMPK binding to WIPI4. **(B)** Phosphorylation of ULK1 at S555**, (C)** pBECN1 at S14, **(D)** TSC2 at S1387. **(E)** Quantification of TSC2 binding to WIPI3. **(F)** Phosphorylation of AKT at S473, and **(G)** ULK1 at S757. All data are shown as relative fold change compared to control condition and were analyzed with an unpaired t-test.; ± SEM; * p < 0.05, **p < 0.01, ***p < 0.001.

**Fig. S5:**
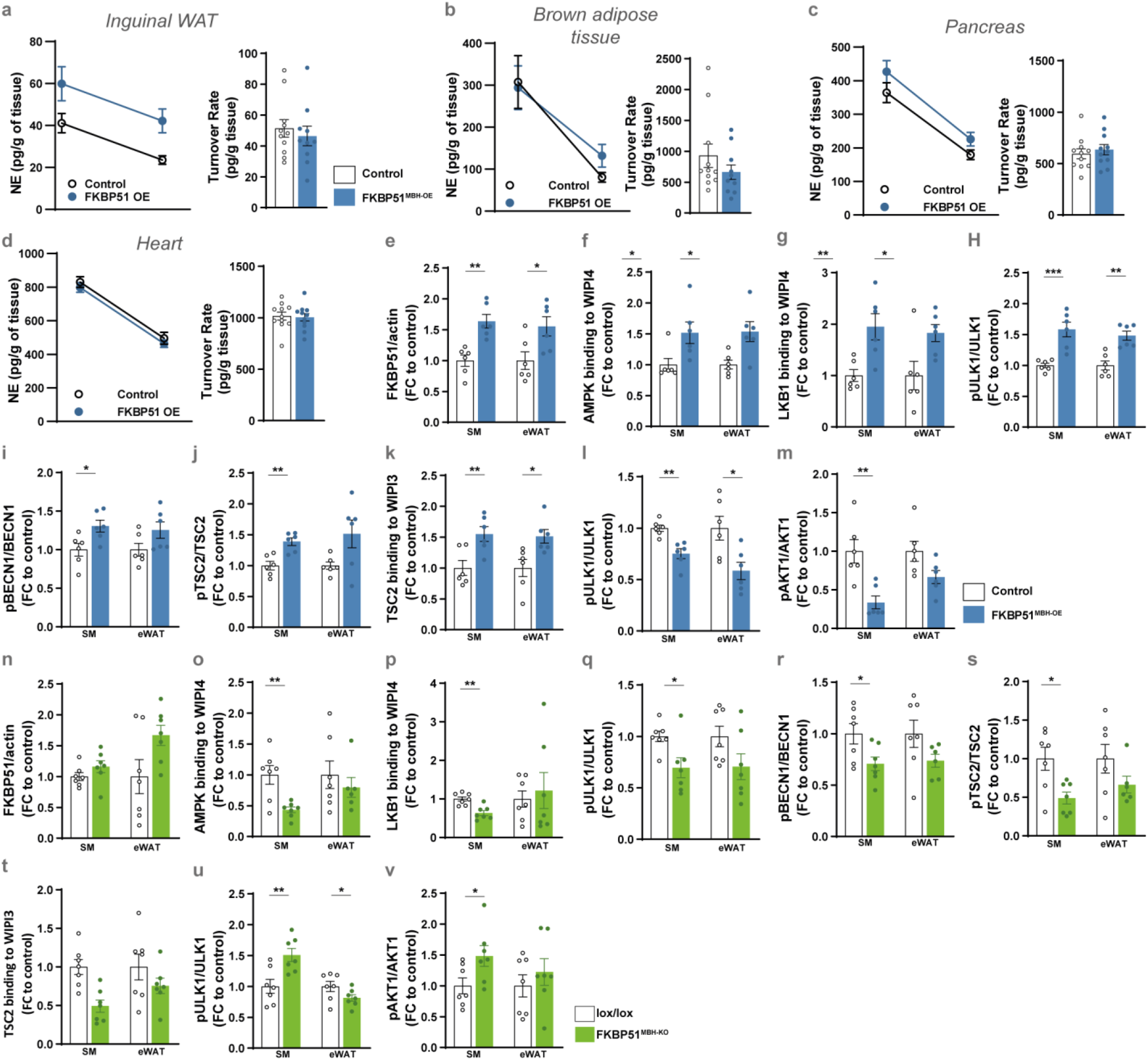
Effects of hypothalamic FKBP51 overexpression on peripheral autophagy signaling. **(A-D)** Representative decrease in tissue NE content after MPT injection (left panel) and turnover rate (right panel) were determined on inguinal WAT and brown adipose tissue (BAT), pancreas, and heart. **(E-V)** Pathway analysis of main autophagy and mTOR marker in the soleus muscle (SM) and epididymal white adipose tissue (eWAT). FKBP51 overexpression is depicted in blue and FKBP51 deletion is depicted in green. **(E)** Quantification of FKBP51 protein level. **(F)** Quantification of AMPK and **(G)** LKB1 binding to WIPI4. **(H)** Phosphorylation of ULK1 at S555**, (I)** pBECN1 at S14, **(J)** TSC2 at S1387. **(K)** Quantification of TSC2 binding to WIPI3. **(L)** Phosphorylation of ULK1 at S757, and **(M)** AKT1 at S473. **(N)** FKBP51 level in SM and eWAT of FKBP51^MBH-KO^ mice. **(O)** Quantification of AMPK and **(P)** LKB1 binding to WIPI4. **(Q)** Phosphorylation of ULK1 at S555**, (R)** pBECN1 at S14, **(S)** TSC2 at S1387. **(T)** Quantification of TSC2 binding to WIPI3. **(U)** Phosphorylation of ULK1 at S757, and **(V)** AKT1 at S473. All data are shown as relative fold change compared to control condition and were analyzed with an unpaired t-test.; ± SEM; * p < 0.05, **p < 0.01, ***p < 0.001.

